# High-throughput cell spheroid production and assembly analysis by microfluidics and deep learning

**DOI:** 10.1101/2022.10.02.510497

**Authors:** Martin Trossbach, Emma Åkerlund, Krzysztof Langer, Brinton Seashore-Ludlow, Haakan N. Joensson

## Abstract

3D cell culture models are an important tool in translational research but have been out of reach for high-throughput screening due to complexity, requirement of large cell numbers and inadequate standardization. Here, we present a high-throughput workflow to produce and characterize the formation of miniaturized spheroids using deep learning. We train a convolutional neural network (CNN) for cell ensemble morphology classification, benchmark it against more conventional image analysis, and characterize spheroid assembly determining optimal surfactant concentrations and incubation times for spheroid production for three cell lines with different spheroid formation properties. Notably, this format is compatible with large-scale spheroid production and screening. The presented workflow and CNN offer a template for large scale minispheroid production and analysis and can be extended and re-trained to characterize morphological responses in spheroids to additives, culture conditions and large drug libraries.

## Introduction

Since 3D cell culture formats can more precisely predict drug response^1,2^, expand the range of *in vitro* culturable cell types and more accurately recapitulate tissue phenotypes unavailable to 2D cell culture^3^, methods and devices for 3D cell culture ranging from complex Organ-on-chip devices^4^ to spheroids^5^ are currently gaining significant momentum.

Spheroids are self-assembled three dimensional clusters of cells that arrange into sphere-like clusters, normally consisting of thousands of cells^6,7^. Spheroids are produced by culturing cells in conditions that promote cell-to-cell contact, while limiting cell attachment to surfaces. Examples of such culture formats include hanging drops^8^, spinner culture^9^, magnetic levitation^10^, low-attachment microwell plates^11^, acoustic manipulation^12^, and microfluidic^13^ formats and have yielded intermediate complexity tissue models representing *e.g* human liver, heart, gut and intestine as well as disease tissue for a wide range of tumors (lung^14^, pancreas^15^, prostate^16^ and ovarian cancers^17^). A collection of published spheroid and miniaturized spheroid models is available in the MiSpheriod database^18^. Here, we refer to these miniaturized spheroids as minispheroids, composed of substantially smaller numbers of cells than conventional spheroids.

While spheroids can provide a more accurate representation of *in vivo* tissue than 2D culture, the use of standard-size spheroids consisting of thousands of cells per spheroid, have been limited to medium-scale high-throughput screening *i.e.*, chiefly limited to tens or hundreds of drug candidates^19^ or drug combinations^20^. Challenges to increasing the scale of screens include the prohibitively high cell material cost^1^, spheroid production capacity and standardization, as well as high-throughput 3D imaging and image analysis. Costs are particularly high for screens using primary material, where *e.g.*, primary hepatocytes typically cost 500$ per million cells. Similar challenges also limit large library combinatorial drug testing in rare cell samples *e.g.*, in functional precision drug screening for individual patients^21^. Minispheroids offer an avenue to overcome these limitations.

Novel microfluidic formats have the promise to truly advance the standardization^22,23^ and throughput^24,25^ of spheroid production by droplet microfluidics to the hundreds of thousands per hour^26^, by a production rate (spheroids produced per hour) definition of throughput. In this context it should be noted that for full screening campaigns, other measures of throughput such as the amount of information per unit time for the entire screening duration can be more informative^27^. Other microfluidics-based spheroid production methods include electrowetting-on-dielectric (EWOD) generated droplets^28^ and microfluidic vessel (MV) based methods^29,30^. Both EWOD and MV have enabled microscale control of the spheroid formation process, but their throughput is limited by chip or well grid surface area.

Droplet microfluidics is a high-throughput platform technology utilizing nano- or picoliter scale aqueous droplets in fluorinated oils stabilized by surfactants^31^ with prominent applications in high-throughput screening^32^, droplet-PCR^33^ and single cell analysis^34^. Fluorinated oils, such as Novec HFE-7500, allow for gas transport to droplets. Biocompatible fluorosurfactants, commonly consist of a hydrophilic polyethylene glycol head group and one or more fluorophilic tail groups^35–37^. Surfactants lower surface tension, stabilize the oil-water interface and ensure low cell adhesion to the droplet interface, which has been key to enabling mammalian cell survival^38^ and proliferation^39^ in droplets.

High cell viability was demonstrated for cells cultured on fluorinated oils with non-ionic PEG-PFPE amphiphilic tri-block copolymer fluorosurfactants^38^ and on Novec HFE-7500 with or without surfactant^23^. Fluorinated oils such as Novec HFE-7500 and Fluorinert FC-40 have also been employed specifically for culture of adherent human cells *e.g.*, keratinocytes^40^ and stem cells^41^, at liquid-liquid interfaces. Nanosheets formed from denatured proteins *e.g.*, albumin, at the interfaces of cell media and Novec-HFE 7500, detected both in the presence of and in the absence of small molecule surfactants in the fluorinated oil, has enabled cell attachment to these surfaces. Notably, the addition of surfactants altering nanosheet mechanics and assembly at the fluorinated oil-aqueous interface, have been demonstrated to impact stem cell adhesion, expansion and phenotype^41^ While small molecule surfactants have been used to enable cell adhesion for liquid-liquid culture, a central purpose of surfactants in droplet microfluidics in general, and cell culture in droplets in particular, has been minimizing cell and biomolecule adhesion.

Aside from the interface, the specific composition of the aqueous phase influences the assembly of cells as well. Droplet microfluidic spheroid production can either use scaffolded, or scaffold-free formats. As scaffolds, a number of gel materials such as alginate^42,43^, collagen^44^, matrigel^45^ or mucins^46^ have been used. Other additives, such as drug cocktails^47^ are also used to promote spheroid formation or cell differentiation. As a result, there is a wide variety of spheroid production formats, incubation times and additives expediting or supporting spheroid formation. In addition to the different responses to these in different cell lines and primary cell samples, rapid and automated methods to survey this parameter space could greatly accelerate spheroid model development, enabling spheroid production from a wider variety of cell types and primary cell samples. Microfluidic formats in general, and droplet-based in particular enable production and handling of spheroids at substantially higher throughput^26,48^ than conventional plate-based formats *e.g.*, hanging-drop culture, or spheroid-forming ULA (ultra low attachment) plates.

In the development of minispheroid production methods as well as in the subsequent analysis of minispheroid screens, most read outs require each minispheroid to be analyzed individually. In spheroid analysis, microscopy imaging provides an information-rich read out and is routinely employed, for larger spheroids^49,50^ as well as for minispheroids^24,25,48^. Datasets of tens to hundreds of spheroids have typically been analyzed manually in software^43,51^ such as ImageJ. To enable large-scale analysis of minispheroids, custom automated scripts^25,45^ in *e.g.*, MATLAB, have been used to determine *e.g.*, spheroid area and circularity. Convolutional neural network based image analysis offers an alternative and complementary route to image analysis to segment, or classify minispheroid images trained from human classification rather than definitions based on area decrease, or circularity.

Deep learning applications range from traffic predictions^52^ over image synthesis^53^ to medical applications^54^ and for example greatly improved the accuracy in diagnostics by assisting the pathologist in image analysis^55–57^. Multilayered artificial neural networks are used for deep learning applications, as they are especially powerful in pattern recognition and the translation equivariance of convolutional neural networks (CNN) makes them the method of choice for tasks pertaining to image data or computer vision^58^. ResNets are deep CNNs with residual connections^59^. These use ungated identity shortcut connections, which allows for flow of information across layers without attenuation^60^ enabling training and use of the layer depth^59^. To perform classification tasks, a network is trained on a manually classified data set. In image recognition, a database widely used for training is ImageNet^61^, consisting of more than 14 million annotated web images of everyday objects. For highly specialized applications *e.g.*, medical image recognition, transfer-learning can be utilized *i.e.*, fine-tuning a pretrained model to perform classification, segmentation or localization on narrower object classes. Multicellular structures in droplets present one set of narrower object classes, where automated and accurate image analysis can have profound impact *e.g.*, effectively enabling multifactorial optimization of a 3D cell culture model.

Since each cell type has distinct self-organization characteristics we have devised a method to enable automatable microfluidic minispheroid generation and imaging as well as an analysis workflow to produce, image and classify standardized minispheroids in a scalable manner (*Figure 1*). We characterized microfluidic cell encapsulation for spheroid production, and increased production capacity. We demonstrate the use of this workflow by applying it to the analysis of spheroid assembly over time at different surfactant conditions for three different cell lines with differing spheroid formation capacities. Cells were encapsulated in 220 μm diameter aqueous droplets in fluorinated oil, using five different surfactant concentrations ranging from 0.5 % to 2 % w/v to study the effect on spheroid formation. The emulsion was transferred into microwell plates and continuously imaged for 15 hours. We then trained CNNs to be able to classify cell ensemble morphologies in droplets and analyze the progression of cell assembly. Taken together, the presented workflow offers an effective and fit-for-purpose approach to increase production and characterization capabilities in microfluidic droplet microtissue engineering. It offers significant scaling potential for future pharmaceuticals screening campaigns in 3D cell models and should enable accelerated minispheroid model development and adoption.

**Figure 1.**
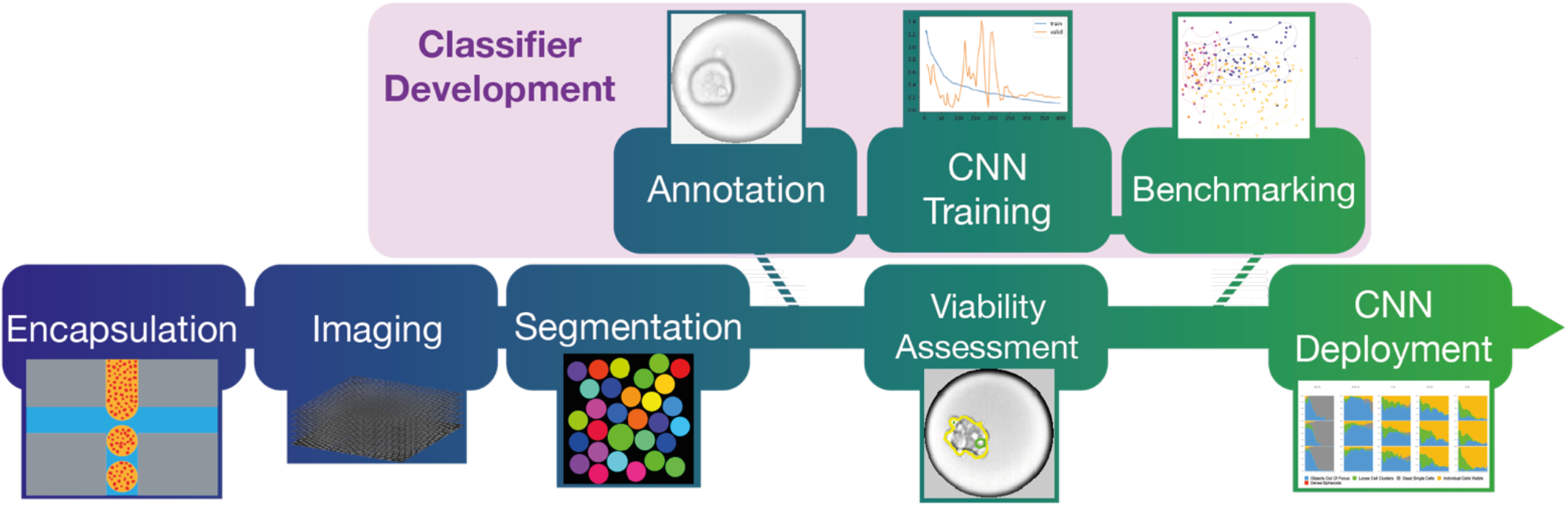
Experimental workflow for CNN categorization. Depicted are the different stages from encapsulation through imaging and network training to analysis. A highly concentrated cell suspension (7×10^6^ cells/mL) is encapsulated in droplets with 240 μm diameter. The emulsion is transferred to an automated microscope and imaged for 15 hours. The resulting images of the monolayer are segmented into single-droplet images. A subset of these images are annotated and used to train a CNN classifier. The classifier’s performance is benchmarked against more conventional means of analysis. Finally, the trained CNN is used to classify the remainder of the images. Additionally, viability of the spheroids at the end of incubation is assessed via fluorescent image analysis.

## Materials & Methods

### Microfluidic Device Fabrication

The negative pressure cell microfluidic devices used were fabricated as previously described^26^. Briefly, a channel design with through holes for inlets and outlets was cut from a 2mm PMMA sheet, according to the channel design (ESI1) to a depth of 100 μm using a CNC (Roland Modela MDX-40). The cut PMMA was sealed with a pressure sensitive adhesive (ARcare). A double-sided adhesive was applied to a 3D printed reservoir and outlet layer (ESI2) and holes were cut in the adhesive using a biopsy punch for inlets and outlets. Finally the full 4 layer device was assembled by combining the 3D printed top part and double sided adhesive with the PMMA channel device.

### Cell Culture

BJ hTert cells were cultured in High Glucose DMEM, 10 % FBS and 100 U/mL penicillin-streptomycin. RT4 cells were cultured in McCoy’s 5a, 10 % FBS, 2mM Glutamine and 100 U/ml penicillin-streptomycin. 5637 cells were cultured in RPMI, 10 % FBS, 2mM Glutamine, 1 mM Sodium Pyruvate, 10 mM Hepes and 100 U/mL penicillin-streptomycin. All cells were cultured in 75 cm^2^ vented cap culture flasks (Corning) in a Sanyo CO2 Incubator at 37 °C and 5% CO_2_.

### Chip and Droplet Generation Characterization

After adding 200 μL of High Glucose DMEM with 10 % FBS and 100 U/mL penicillin-streptomycin to the aqueous reservoir, we generated droplets in a continuous phase of Novec HFE-7500 (3M) with 1 % w/v 008 surfactant (Ran Biotechnologies, Boston, USA), a non-ionic PEG-PFPE amphiphilic tri-block copolymer fluorosurfactant, at −2.5, −5, −7.5, −10, −12.5, −15 and −20 kPa. We recorded the lapsed time between engaging the pressure controller and the appearance of air bubbles stemming from the aqueous reservoir. After that, we measured any residual volume in the reservoir that failed to enter the chip. Droplets were imaged on a Nikon TI-Eclipse, using 10x magnification. We manually measured the diameter of the droplets using FIJI. Using static electricity with a Zerostat 3 (Milty)^62^ we unified and removed the aqueous phase and measured the total collected oil volume by pipetting.

### Droplet Microfluidic High-Throughput Spheroid Generation

BJ hTert, RT4 and 5637 cells were cultured to 80-90 % confluence and harvested using TrypLE Express (Gibco). Cell concentrations were determined with a TC20 Automated Cell Counter (Bio-Rad) and cells were resuspended at ~ 7.5×10^7^ cells/mL in their respective media. Prior to encapsulation the cells were stained with POPO-1 iodide (0.05 μM), Green Caspase3/7 (0.25 μM), Image-iTTM Red Hypoxia Reagent (5 μM) and Cell Tracker DeepRed (0.1 μM) (all from Invitrogen). Cell samples were split into five equal volumes and subsequently encapsulated in 5.6 nL droplets in a continuous phase of fluorinated oil Novec HFE-7500 (3M) with 0.5 %, 0.75 %, 1 %, 1.5 % or 2 % w/v 008 surfactant (Ran Biotechnologies, Boston, USA), respectively. The emulsion was generated by applying −12 kPa pressure, generated by a PC20 pressure controller (NAGANO KEIKI CO., LTD., Japan) at the collection outlet of the microfluidic cell encapsulation device.

### Droplet Imaging

RT4 and 5637 cell samples were transferred to 3 wells per sample and BJ hTert to 5 wells per sample in a 96 well plate (Perkin-Elmer). The wells were sealed with a gas permeable sealing film (Axygen) and imaged in bright-field mode with a 10x objective every hour for 15 hours on an Opera Phenix HCS instrument (Perkin-Elmer) at 37 °C and 5 % CO2. Images were captured in a stack of 2-7 layers to ensure an adequate focal plane for each well, each containing 9 tiles, to cover the entire microtiter plate well. After 15 hours droplet fluorescence imaging at 488/522 (Caspase3-7 green), 425/456 (POPO-1 iodide), 640/706 (CellTracker) and 490/610 (Hypoxia) was performed.

### Segmentation and CNN Training

The scripts developed here can be found in the supplementary information. The bright-field image data sets were exported and each droplet region of interest segmented using a MATLAB (MathWorks) script based on the Hough transform, for a total of 16956 single-droplet images. A subset, consisting of 856 images (95 Centered Aggregates, 87 Centered Spheroids, 50 Single Cells, 155 Non-Centered Aggregates, 86 Non-Centered Spheroids, 101 Out Of Focus Objects, 282 Faulty Pick Ups) of cropped, single-droplet images were manually classified as one of the 7 categories. The image set was used to train a convolutional neural network (CNN) using the fastAI library^63^. Prior to training the SpheroClass-CNN, the labeled data set was resized to 128×128 pixels and split into a training set containing 80 % of the images and a validation set containing the remaining 20 %. These image sets were expanded through data augmentation such as zooming, warping and rotating. We used a ResNet50, deep residual network architecture consisting of 49 convolutional and one fully connected layer with ReLU functions as non-linearity, pretrained on ImageNet^59^. The base learning rate was determined using the fastAI library’s Learning Rate Finder, followed by Automated Learning Rate Suggester^64^. The batch size was set to 64 images. The SpheroClass CNN was trained for 300 epochs and through a callback function the model iteration with the lowest error was retained. A second CNN was similarly trained to distinguish between the two out-of-focus categories.

### Benchmarking of Cell Encapsulation and CNN

Cell viability before encapsulation was assessed using trypan blue and an automated cell counter (Biorad). Fluorescent images of the droplets were captured in the automated imager at the end of the 15 h incubation period. These fluorescent images were segmented into single-droplet images by an adapted version of the bright-field image segmentation script in MATLAB (see supplementary information). The cells in RT4 and BJ hTert sample images were segmented using fluorescent signal from the CellTracker dye. 5637 cells’ observable cell area was segmented by hand using a tablet and stylus pen (Apple). Finally, the cell area was compared to POPO-1 iodide-positive area, which was defined as pixels with a signal intensity higher than 10000 out of 65435.

To benchmark the CNN, we manually segmented the cell-occupied area of 231 individual droplet images containing RT4 cells of 0.5 %, 1 % and 2 % w/v surfactant samples taken at various time points with a tablet and stylus pen (Apple) and extracted the segmented objects’ area, circularity, brightfield inner contrast and centroid in relation to the droplet centroid using MATLAB scripts. The resulting data was then compared to the CNN’s prior classification.

### Morphological Classification

The trained model was used to classify the images of the data set. 16221 single droplet images were analyzed with the SpheroClass CNN and classified as one of the following categories: “Faulty Pickup”, “Out Of Focus Object”, “Single Cells”, “Non-Centered Aggregate”, “Centered Aggregate”, “Non-Centered Spheroid” or “Centered Spheroid”. In a subsequent round, a second model was used to sub-classify the images labeled “Out Of Focus Object” as either “Out Of Focus Non-Spheroid” or “Out Of Focus Spheroid”. Classifications in either model that did not meet the minimum probability of 0.75 assigned to the prediction were discarded.

## Results & Discussion

### Microfluidic spheroid production throughput characterization

In order to improve minispheroid production rate we characterized compressor and pressure controller actuated microfluidic droplet production capacity, and compared this with liquid handling robot actuation. We characterized flow rates and droplet generation rate as well as aqueous to fluorinated oil ratio and droplet size of the resulting emulsion in a negative pressure range between −2.5 and −20 kPa. At −2.5 kPa the chip failed to produce any droplets and instead only oil was collected. At applied pressures below −2.5 kPa the microfluidic device produced droplets (*Figure 2*). Droplet size decreased with increasing pressure differential, plateauing above −12.5 kPa.

**Figure 2.**
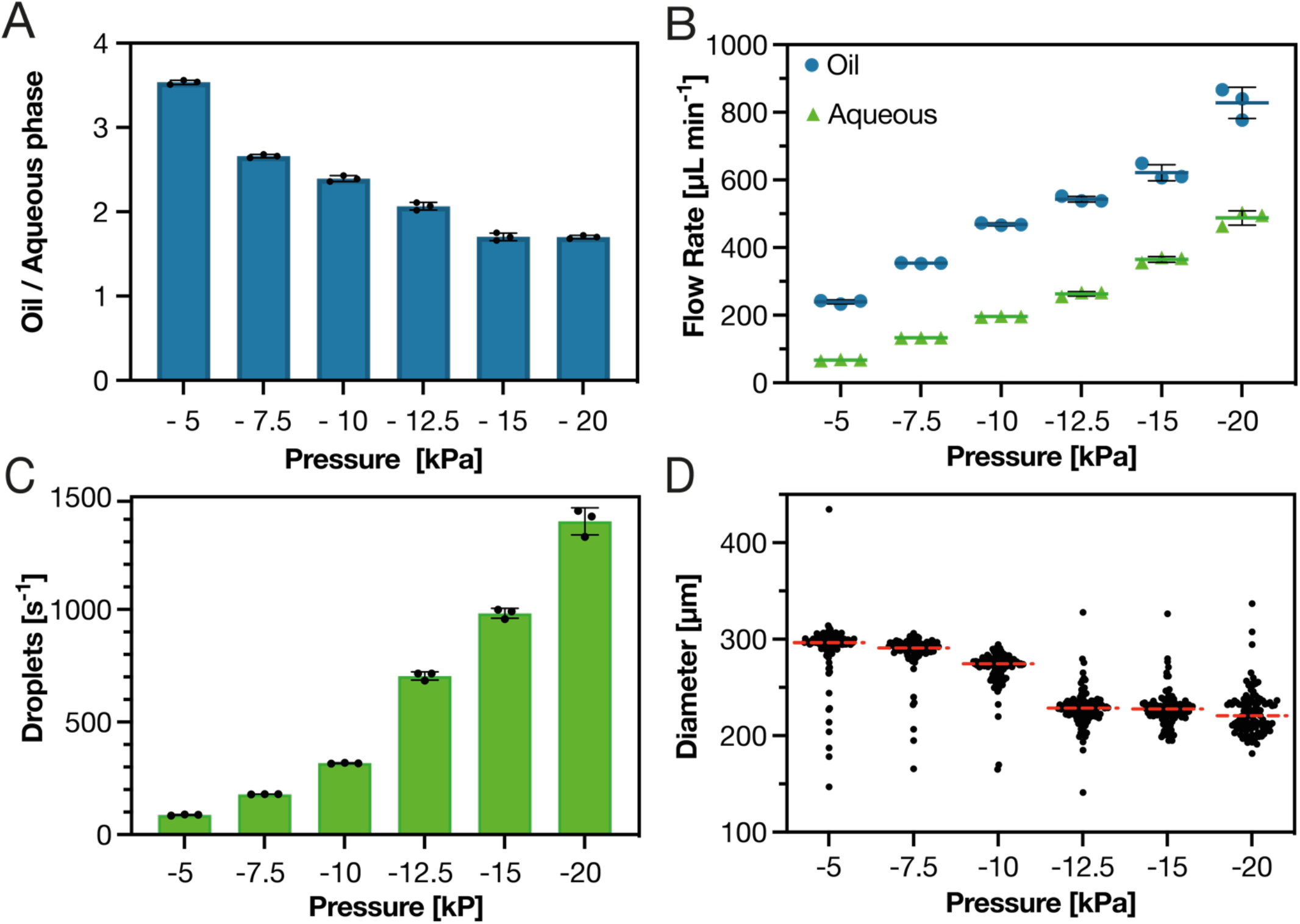
Negative pressure droplet generation. A) Bar plot showing the volume fraction of fluorinated oil normalized to the volume of the unified aqueous phase (three technical replicates, black lines signify one standard deviation). B) The flow rates of the oil and aqueous phase, respectively, as a function of the applied pressure differential (three technical replicates, colored lines illustrate means and black lines one standard deviation). The dependency of droplet generation rate is shown in C) and the diameter of the droplets generated in D) (red, dashed lines illustrate the mean, n=671), with varying negative pressures applied at the outlet. With −2.5 kPa outlet pressure only oil and no droplets were collected.

Droplet production rate increased with increasing pressure differential, resulting from the decreasing droplet sizes as well as increasing flow rates.

Notably, the volume fraction of aqueous phase in the collection increased as a function of the pressure, leading to lower oil consumption per generated droplet with increasing pressure differential. Compared to previously published figures for negative-pressure, pipette-actuated droplet production in similar devices, pressure controller actuated droplet production at −15 kPa increased production rate more than 20-fold. Summarily, the chip showed good robustness over a considerable range of pressures with high inter-experimental reproducibility.

### Encapsulation, imaging and segmentation

In order to characterize cell assembly and spheroid formation in microfluidic droplets, cells were encapsulated at −12 kPa to obtain an emulsion of droplets with a diameter of 220 μm (~5.6 nL). The emulsion was transferred to a 96-well plate. Imaging and subsequent segmentation generated cropped images, each containing an individual droplet. In this way we ensured a consistent focus for the morphological analysis of the droplet contents (*Figure 3A)*. The size distribution of the segmented droplets was sufficiently narrow with a wide enough safety margin to the gating limits to ensure unbiased segmentation, with a mean diameter of 218 μm and a *CV* of 8.3 % (*Figure 3B)*. This suggests high repeatability as well as absence of pronounced shrinkage due to osmotic pressures differences or evaporation.

**Figure 3.**
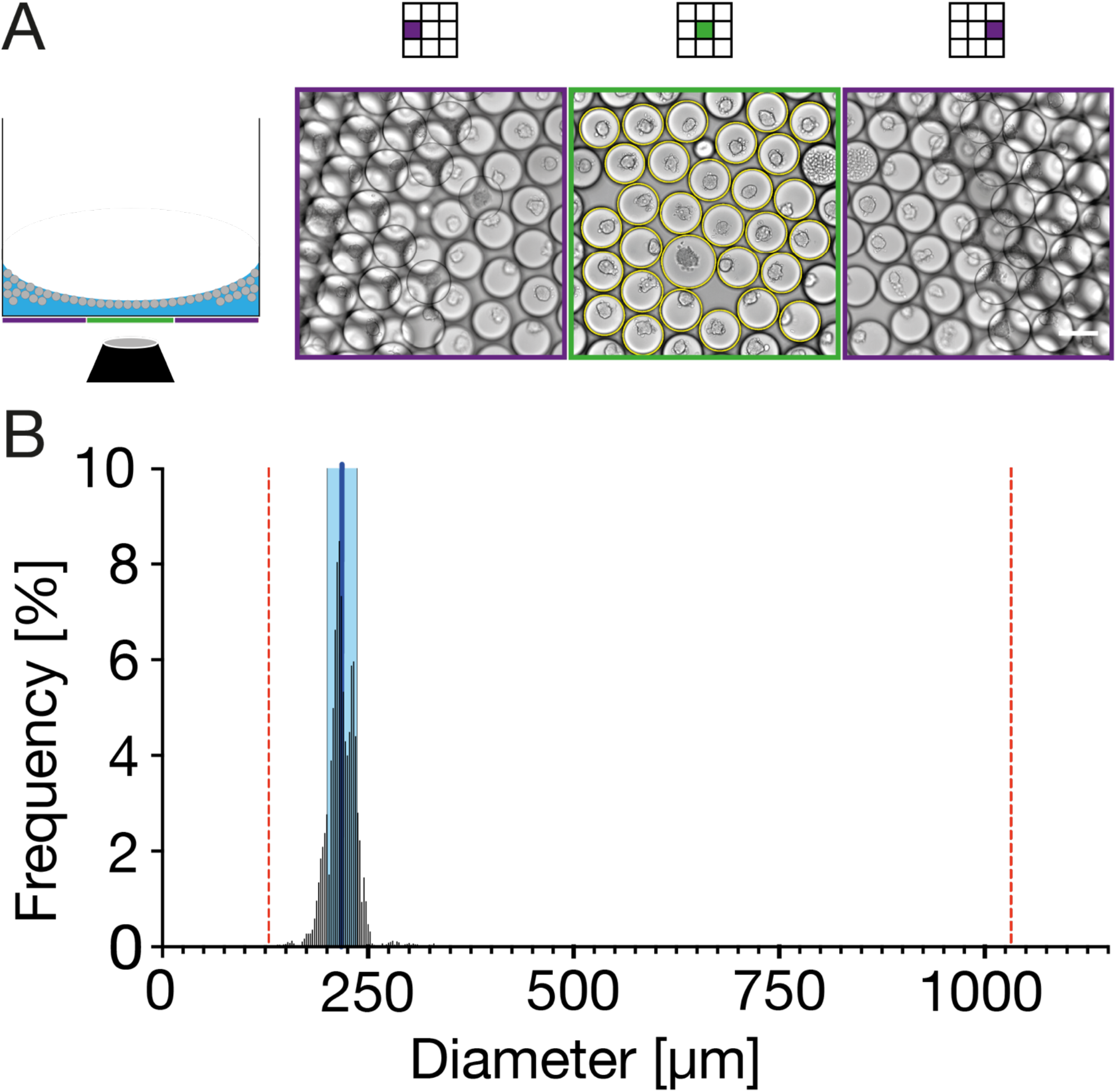
Droplet imaging and segmentation. A) Imaging illustration. Left: 2D-schematic of well plate imaging in the automated microscope. The oil (blue) forms a meniscus in the well, leading to a monolayer of droplets (gray circles) at the lowest point of the oil meniscus and multilayers at the periphery. Right: Bright-field images. Green square shows the center tile with yellow circles highlighting the segmented droplets subsequently fed to the CNN for categorization, the purple squares show two periphery tiles (white scale bar = 200 μm). B) Histogram showing the size distribution of segmented droplets. The light blue box illustrates standard deviation, the dark blue line the median size. Red broken lines mark the minimum and maximum allowed apparent diameter, respectively. The emulsion stays monodisperse throughout the duration of the experiment (n = 16956, 0.5 μm binning).

Due to gravity, all cellular droplet contents should come to rest in the same focal plane *i.e.*, the lowest point of the droplet. Any deviation from that suggests additional mechanisms affecting the cells, such as fluidic microcurrents, cellular interaction with the interface or changes in cell buoyancy, causing the cells to leave the focal plane.

Indeed, cell assemblies displaced from the bottom of the droplet were observed in some instances (*Figure 4*). Since the droplets were not agitated, we can exclude micro- or macrocurrents as the cause for this phenomenon (see also *Supplemental Figure 1*). The consistency with which cell assemblies consistently orient at the bottom center of the droplets for many of the cases tested demonstrate that such agitation was not present in the wells during incubation. Thus cell interaction with the water-oil-interface is likely, possibly mediated by the protein nanosheets, which has been demonstrated to form at the interfaces between BSA or FBS containing aqueous media^65^.

**Figure 4.**
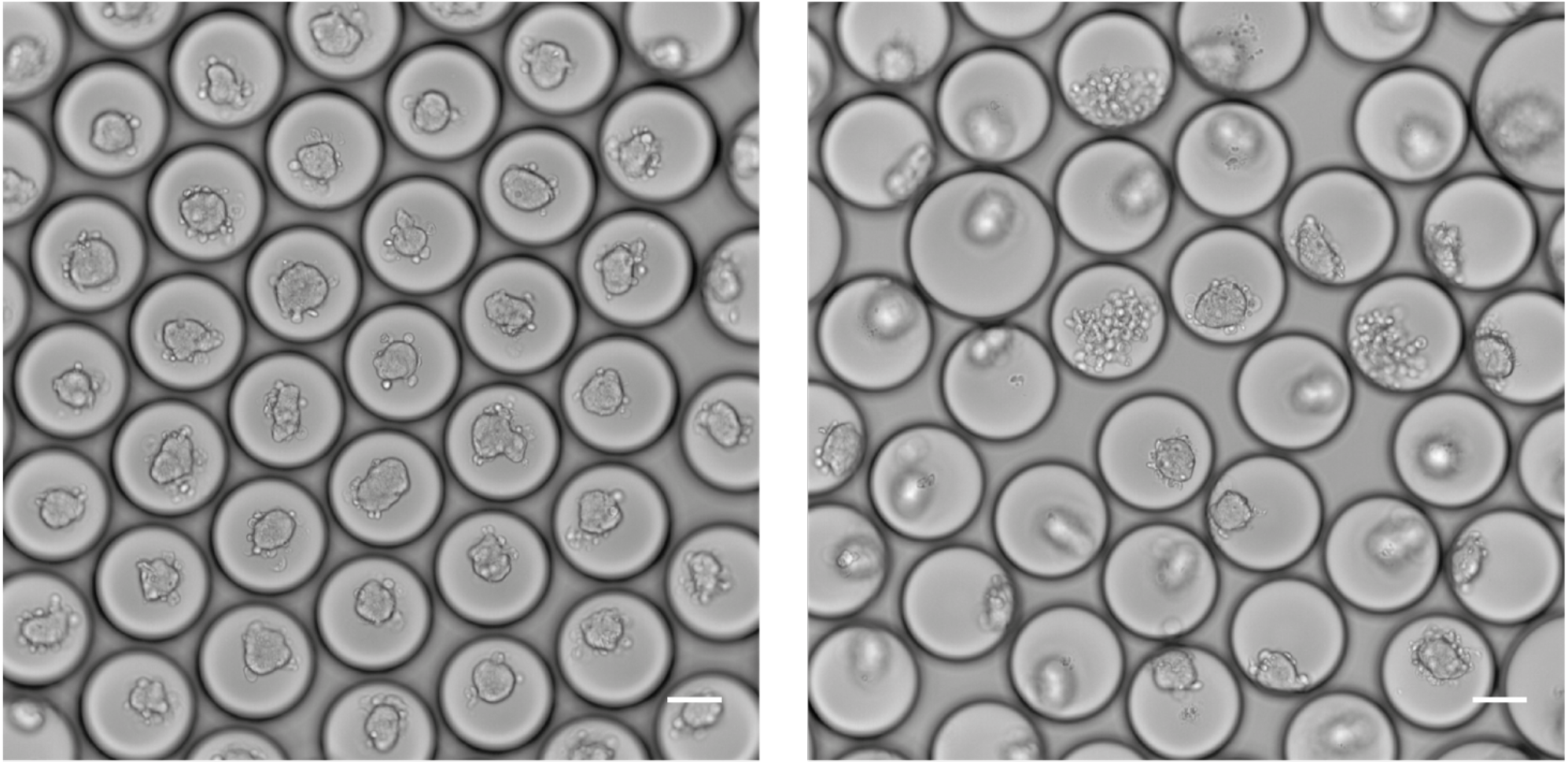
Images of droplet monolayer. The images show RT4 cells after 13 h of droplet cell culture. In the left image the cell assemblies are all in the same focal plane and approximately laterally centered *i.e.*, close to the lowest point of the droplet. In the right image however, most cell assemblies are out of focus and removed from the lateral center point of the droplet (white scale bar = 100 μm).

Consistent with established spheroid generation methods, minimizing interaction with the available interface results in faster and more robust spheroid assembly; hanging drop techniques utilize a water-air interface and culture in U-bottom plates generally employs specially treated ultra low attachment surfaces.

We hypothesize that both laterally non-centered and out of focus position of ensembles indicates interaction between cells and interface, in concordance with what has been previously described^41^. To further characterize this, we segmented a total of 16956 single droplet images (5552 images for RT4 cell line, 6008 images for BJ hTert cell line, 5396 images for 5637 cell line), resized to 128×128 pixels for morphological classification. To validate the droplet segmentation method, 9 representative images were selected, and subjected to both automated and manual droplet segmentation. In these images, 99% of manually identified droplets were successfully segmented by the MATLAB script (*Supplemental Figure 2*).

### Training and deploying the CNN

From the 16956 images, seven distinct cell ensemble morphology categories along with a category for erroneously segmented droplets, termed Faulty Pick Up, were identified. A subset, consisting of 856 of the cropped, single-droplet images were labeled with one of the 7 categories to train the CNN. The learning rate is an important hyperparameter to tune for a CNN. Based on Learning rate finder and Suggester functions, a learning rate of 9.12 × 10^−3^ was used (see *Supplemental Figure 3A*).

The augmentation, which presented the network with a different data set for each training period, or epoch, in combination with the residual network already being pre-trained on ImageNet, enabled the training of a CNN despite the modest size of the spheroid data set. The SpheroClass-CNN reached top performance after 102 epochs of training with a 0 % error rate during validation.To verify that the trained model would perform as expected, we applied the SpheroClass-CNN to a small set of images that had not been part of the training or validation set. The results were compared to the images’ manual annotation (*Figure 5*).

**Figure 5.**
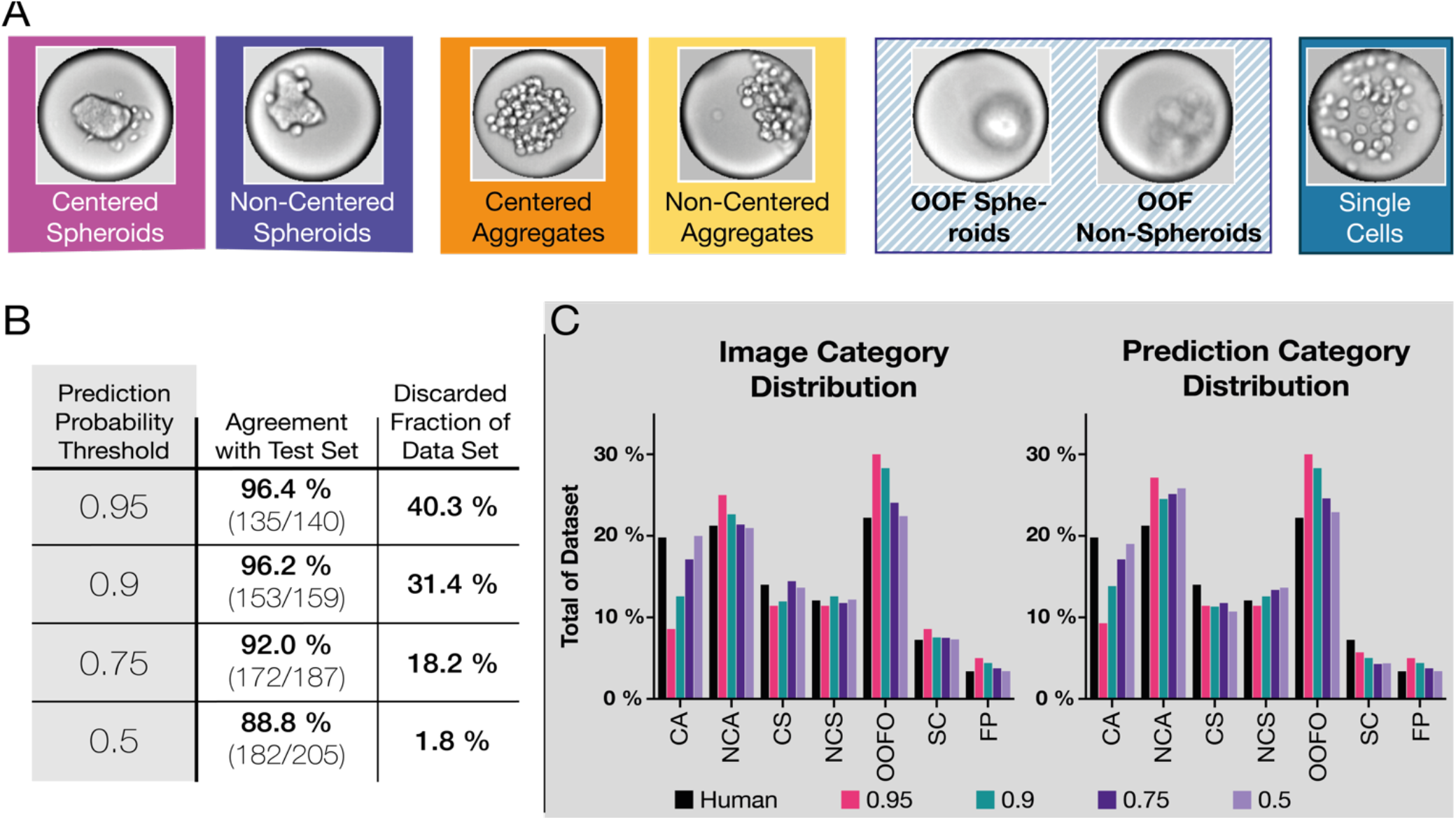
SpheroClassCNN performance results guiding thresholding considerations. A) Examples of morphological categories, differentiating between 3 different overarching morphologies (Single Cell, Aggregate and Spheroid) as well as location within the droplet (Centered, Non-Centered, Out Of Focus). B) Table illustrating the trade-off between accuracy in the validation and preservation of data points. While lowering the prediction probability threshold from 0.75 to 0.5 leads to a decrease in size of the discarded data set by an order of magnitude, a mere 10 out of the 18 additional droplets (55.6 %) were correctly classified in the test set (test set n = 207, data set n = 16956). C) Bar plots showing the category distribution of analyzed images (left) and the resulting predictions (right) using different probability thresholds. Very stringent thresholds introduce considerable selection bias, most notably in the Centered Aggregate and Out Of Focus Objects categories.

Following evaluation of the test set, we set the probability threshold of the model’s output at 0.75, meaning that the model assigned at least 75 % probability to the respective image’s most likely classification, and analyzed the remainder of the images, disregarding the predictions that did not meet this criterion. We chose this threshold considering the apparent trade-offs between accuracy and resulting data set size, and limitation of selection bias (*Figure 5C*). Using the 0.75 probability threshold yielded 13263 data points from originally 16221 single-droplet images, representing a loss of 18.2 %. The certainty *i.e.*, assigned probabilities, with which images containing Centered Aggregate spheroids could be identified as such, was lower than that for images of other category objects (see *Supplemental Figures 4* & *5)*. This trend was observable in the test set as well. Assessment of confusion matrices over different probability thresholds supported this notion further: Aggregates, centered or non-centered, comprise arguably the most diverse morphological group, so less certainty upon classification is to be expected (*Supplemental Figure 6*).

Analyzing the results of the SpheroClassCNN classification, we noticed that the Out Of Focus (OOF) category made up a large fraction of the images (see *Supplemental Figure 7*). Hypothesizing that we could further resolve this, we employed a second CNN, similarly trained, to distinguish between two distinct out-of-focus object variants. We manually annotated 100 images each for Spheroids Out Of Focus or Non-Spheroids Out Of Focus respectively, by checking the actual morphology in a different focal plane than the one analyzed, and trained the BlurredObjectsCNN using a ResNet50 over 50 epochs with a base learning rate of 1.58 × 10^−2^, achieving 0 % error during validation after 14 epochs (see *Supplemental Figure 3B*).

### Benchmarking of Cell Encapsulation and CNN Classification

Cell viability in microfluidic droplets for spheroid assembly has been demonstrated previously^24,26,46^. Results from a similar microfluidic device, oil-surfactant system and incubation times yielded a 4 % viability decrease from 96.2 % following spheroid retrieval^26^ which is expected to progressively deteriorate with long term incubation *i.e.*, droplet culture for more than 24 h (see *Supplemental Figure 8*). With this and the relatively short incubation period in mind, we chose to monitor cell viability as a quality control measure by including a dead stain, POPO-1 iodide, in the encapsulation media. The viability of 5637 cells was measured to be 99 % before encapsulation; for RT4 and BJ hTert cells, viability was 98 % and 96 %, respectively. To not induce phototoxic effects confounding our analyses we did not fluorescently image the spheroids continuously, but limited exposure to fluorescent light to endpoint imaging after 15 hours to monitor compatibility of the cells with the incubation format. The dead stain, POPO-1 iodide, binds to nucleic acids of cells with compromised outer membranes and thus does not stain the whole interior of the cell. This adds to the inherent difficulties assessing the viability of a 3D cell ensemble with 2D images. Consequently, analysis of the stained area in these experiments is to be considered qualitative. In our analysis we calculated stained area over observable cell area for each droplet (see *Figure 6A*) and concluded that encapsulation and subsequent incubation in droplets does not affect viability negatively: The samples encapsulated in HFE with 1 % w/v surfactant maintained, by our measures, a mean viability of 99.5 % (5637) and 97.3 % (RT4). BJ hTert showed the lowest mean viability with 92.6 %, which is still within reason and might be improved by reducing droplet incubation time (*Figure 6B*). 65.6 % of droplets containing 5637 cells showed no POPO-1 iodide signal above the threshold at all, which we consider to be 100 % viability. We observed fewer droplets with 100 % viability with RT4 (18.7 %) and BJ hTert (9.2 %). A comprehensive graph showing the viability of cells for all conditions can be found in the supplementary material (*Supplemental Figure 9*).

**Figure 6.**
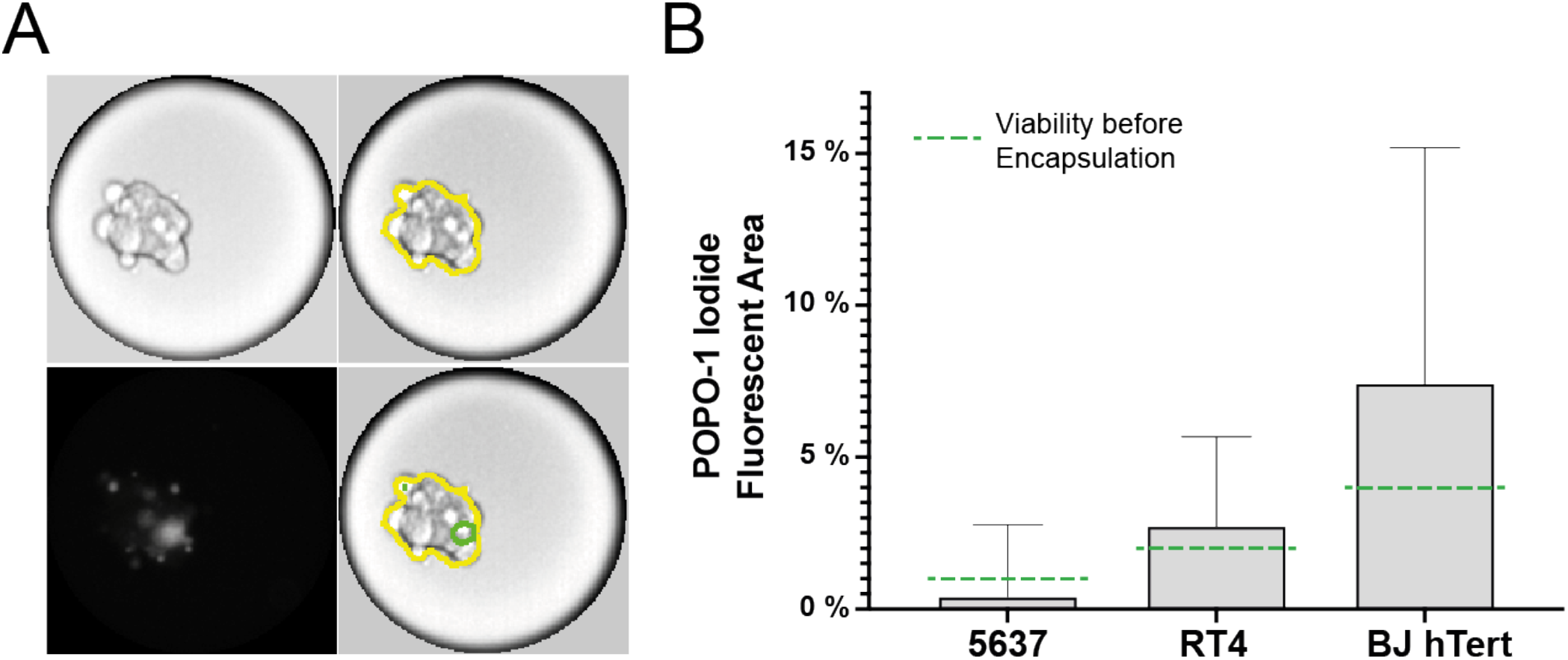
Popo-1 iodide fluorescence analysis after 15 hours. A) illustrates segmented cell area (yellow) and Popo-1 iodide-positive area (green) from the bright field and fluorescence image on the left. B) Plot showing the results of the POPO-1 iodide fluorescent area analysis. Gray boxes illustrate the mean for the respective cell lines in the 1 % w/v surfactant samples, whiskers represent the standard deviation. Green bars signify the viability of single cells before encapsulation.

To assess the CNN’s classification of the cell ensembles, we benchmarked the output from the classifier against conventional image analysis. When researchers assess spheroid production quality, focus is often on uniformity in compactness, sphericity and size^66^. Spheroids can be categorized by these proxy measures or by manual inspection of cell ensemble images. To be able to evaluate the CNN’s performance against conventional image analysis, we manually segmented the cells in 231 images of droplets containing RT4 cells. From these segmented cell area images we measured area, circularity, centroid position in relation to droplet center and internal contrast (see *Figure 7*). In aggregate, these measurements show that as the cells aggregate, the segmented cell area decreases. Circularity increases between 0 h and 9 h of incubation as the cells form ensembles. However, at 14 h after encapsulation the increase in mean circularity had stopped. The mean inner contrast of the segmented cell area reduces over the course of the investigated time points. The decrease in inner contrast is one detectable proxy measure of the condensation of cells in addition to production of extracellular matrix, rendering individual cells less and less discernible.

**Figure 7.**
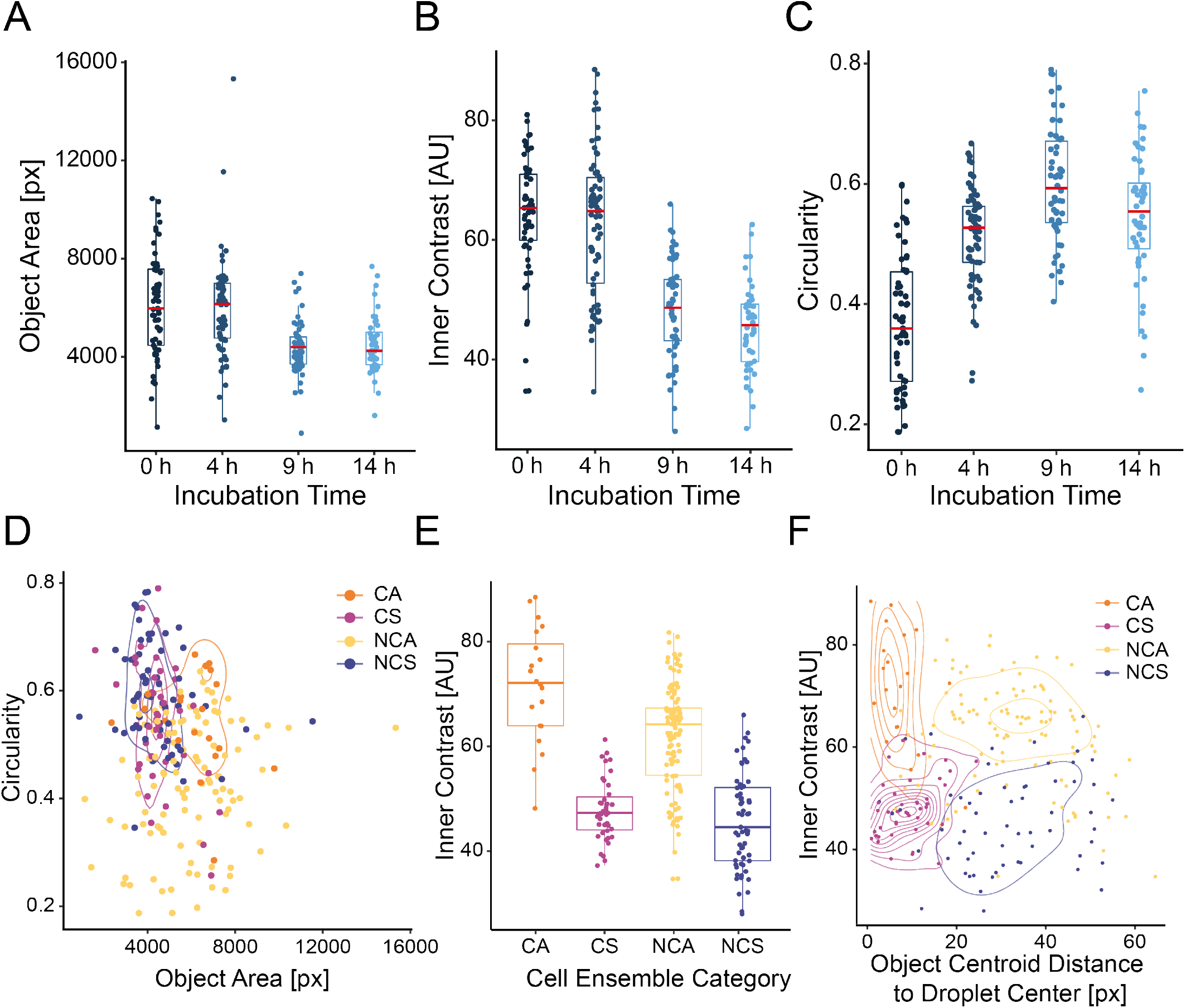
Benchmarking of CNN classification against conventional spheroid assessment parameters on RT4 cell images. A), B) and C) show object area, inner contrast and circularity, respectively, over time for a set of 231 images manually segmented for cell area. Area and inner contrast decrease over the course of imaging, whereas circularity increases with a peak recorded mean value at 9 h incubation time. D) plot shows the CNN classified objects’ circularity over object area. Aggregates’ circularity and area span a wide range, consistent with the diverse appearance of these objects. E) shows the respective inner contrasts for the different CNN classified groups. Objects classified as spheroids by the CNN consistently exhibit a lower inner contrast compared to CNN classified aggregates. F) illustrates the segmented cell objects’ inner contrast over the distance to the droplet center color coded by CNN assigned classification with overlaid color-coded density mapping.

Comparing the respective circularities of the visually distinct categories Centered Aggregate, Centered Spheroid, Non-Centered Aggregate and Non-Centered Spheroid, does not enable discrimination between the categories. Taken together with area, spheroidal categorizations group a little closer together, but are not distinct enough to gate for them. The cell ensembles in the Non-Centered Aggregate category were on average less circular than other categories and had the most varied morphology with regards to circularity and area. Thus, circularity on its own is a poor proxy of overall morphology and does not enable differentiation between different phenotypes. However, intra-object contrast largely enables discrimination between aggregates and spheroids. Plotting the inner contrast over the distance of the segmented cell mass’ centroid to the center of the droplet enhances grouping of the different CNN-predicted classes, but does not segregate them completely.

### Analysis of classification output

Our two-tiered approach categorized 735 segmented images (representing 4.34 % of the data set) as Faulty Pick Ups, which were disregarded for the analysis. The remaining images revealed that the three examined cell lines reacted differently to the varying surfactant concentrations and incubation time (*Figure 8*). 5637 cells did not yield appreciable spheroids but instead less robust cell clusters, even at the highest surfactant concentration. In the 0.5 % surfactant concentration sample however, initial low-density cell clusters disintegrated into single cells, a behavior that was not observed appreciably at the 0.75 % or higher surfactant concentration samples and not at all with the other cell lines. BJ hTert cells showed the most robust spheroid assembly. In all concentrations a majority of the droplets contained centered spheroids after some time had passed; increasing surfactant concentration seemed to merely accelerate this process: After 8 hours, 80.00 % of droplets contained centered spheroids in the 0.5 % surfactant sample, whereas in 85.29 % of droplets centered spheroids had assembled in the 2 % surfactant sample after only 3 hours.

**Figure 8.**
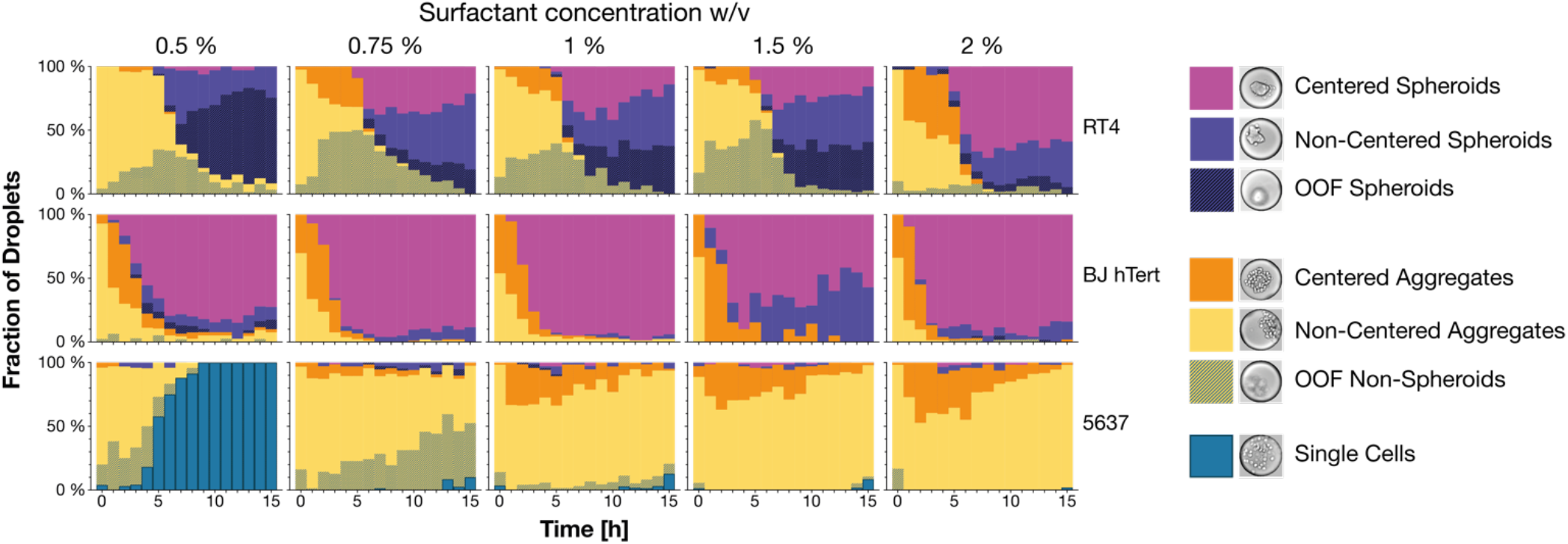
CNN classification of droplets. The graphs show five different surfactant concentrations for the three examined cell lines, RT4, BJ hTert and 5637. With increasing surfactant concentration RT4 cell ensembles exhibit decreasing out of focus behavior. In BJ hTert cell samples, increasing surfactant concentration appears to impact the morphology of the cell ensembles less than with the other cell lines, but accelerate the spheroid formation. 80 % of droplets contain centered spheroids in the 0.5 % surfactant sample after 8 hours, compared to 85 % in the 2 % surfactant sample after 3 hours. 5637 cells display a distinctive single-cell morphology in the lowest surfactant concentration sample, which all but disappears in the higher surfactant concentrations (n=16221). OOF = Out Of Focus.

Finally, RT4 cells formed spheroids at a comparable rate as BJ hTert. The RT4 cells were, however, more prone to interface interactions, as only at the highest examined surfactant concentration a majority of the droplets contained centered spheroids (58.93 %) and very few out-of-focus spheroids (5.36 %). The samples with 0.5 % surfactant concentration on the other hand contained 91.8 % spheroids that were out of focus (67.21 %) or non-centered (24.59 %).

When we break down the results according to centered versus not-centered objects the same general trends are visible in the overall results: both the individual cell line’s capacity to form spheroids and the surfactant concentration correlate positively with the centeredness of droplet contents (see *Supplemental Figure 10*), decreasing out-of-focus behavior. We note that increases in the prevalence of spheroid formation from cell ensembles correlates with increased surfactant concentration for the two cell types that formed spheroids in the experiment. In addition, prevalence in spheroid formation also correlates with increases in cell ensemble centeredness and remaining in focus. Given the aim of rapid and standardized minispheroid formation, the CNN model enables discrimination of these desirable cell states and can therefore be used to guide decisions on incubation time and surfactant concentration used, and could provide guidance on additional parameters *e.g.*, pharmaceutical or media additives which speed spheroid formation. The specific mechanism behind the cells’ interaction with the droplet interface is hypothesized, based on prior studies with the same fluorinated oils, to be promoted by the formation of proteinaceous nanosheets^41^ In studies without surfactants or with other surfactants, these attachments have been implicated in transcriptome alterations in stem cells. As the cell ensemble movement to non-centered and out-of-focus cell ensembles is interpreted as interaction with the droplet interface, it is conceivable that non-centered and out-of-focus cell ensembles might display equivalent discrepancies when compared to centered ones.

Improving process control of cell spheroid production is of great significance to the field, which is limited by the lack of reproducibility and standardization^67^. Enhancing characterization capabilities during production is a prerequisite for overcoming these obstacles. As each compartmentalized droplet is a technical replicate, the statistical power of our method is multiple times better per screened well than a conventional well plate format. The high-throughput nature of our setup could be applied to enable the study of *e.g.*, nano-architectonics^59,60^ and mechanosensing^68^ in stem cell differentiation on a larger scale. Furthermore, entire drug screens could be transferred into this microfluidic droplet format and analyzed with a trained CNN.

## Conclusions

Here, we presented a deep-learning supported approach to high-throughput microtissue production and characterization. We used it to optimize efficiency at the beginning of a 3D cell culture model development. Based on our results, we can determine the appropriate amount of surfactant for any desired spheroid or aggregate morphology, making the whole process more sustainable, cost-effective and establishing the minimal time-in-droplet, both improving throughput and reducing exposure of the cells to a nutrient-limited environment. BJ hTert cells for example do not seem to require more than 0.75 % w/v surfactant to reliably and rapidly form spheroids, but the shortening of assembly time observed in higher surfactant concentration samples might still be worthwhile in specific applications. Utilizing bright-field imaging reduces experimental complexity and minimizes artifacts caused by staining as well as phototoxic effects. Fluorescence imaging can nevertheless be added to complement the data, as was done in the current study.

The use of a trained artificial intelligence algorithm, although limited to a comparatively small training data set, increased the throughput drastically compared to manual annotation of the whole data set with good accuracy. As such, it offers high-throughput morphological characterization for spheroids and is not only applicable to spheroid production but could also be used for drug screening workflows. With increasing annotated data set size, the accuracy should improve further and even more granular analyses of phenotypes should be achievable. Moreover, a CNN trained in such a fashion could be used to study any morphological phenomena of interest on a larger scale, for example in a high-throughput study of nano-architectonics in stem cell research^41^, and an adapted workflow could enable scaling up of stem cell differentiation^40^. Ultimately, the presented study offers a step towards improved process repeatability and robustness by defining and controlling the end product, cell spheroids.

## Author Contributions

**Martin Trossbach:** Conceptualization, Investigation (lead), Software, Data curation, Visualization (lead), Writing-Original draft preparation, Reviewing and editing. **Emma Åkerlund:** Conceptualization, Investigation (support), Writing-Reviewing. **Krzysztof Langer:** Conceptualization, Investigation (support), Writing-Reviewing. **Brinton Seashore-Ludlow:** Supervision, Writing-Reviewing. **Haakan N. Joensson:** Supervision, Conceptualization, Visualization (support), Writing-Original draft preparation, Reviewing and editing.

## Acknowledgements

This work was funded by the *VINNOVA project “Sweden - a test bed for future nanoscale drug development“* and the *Knut and Alice Wallenberg Foundation*. The authors would like to thank Marta Garnelo for her insights during fruitful discussions regarding neural networks.

