## Supplementary Material for "High-throughput cell spheroid production and assembly analysis by microfluidics and deep learning"

#### ResNet50 Configuration

```
ResNet(
  (conv1): Conv2d(3, 64, kernel_size=(7, 7), stride=(2, 2), padding=(3, 3), bias=False)
  (bn1): BatchNorm2d(64, eps=1e-05, momentum=0.1, affine=True, track_running_stats=True)
  (relu): ReLU(inplace=True)
  (maxpool): MaxPool2d(kernel_size=3, stride=2, padding=1, dilation=1, ceil_mode=False)
  (layer1): Sequential(
    (0): Bottleneck(
      (conv1): Conv2d(64, 64, kernel_size=(1, 1), stride=(1, 1), bias=False)
      (bn1): BatchNorm2d(64, eps=1e-05, momentum=0.1, affine=True, track_running_stats=True)
      (conv2): Conv2d(64, 64, kernel_size=(3, 3), stride=(1, 1), padding=(1, 1), bias=False)
      (bn2): BatchNorm2d(64, eps=1e-05, momentum=0.1, affine=True, track_running_stats=True)
      (conv3): Conv2d(64, 256, kernel_size=(1, 1), stride=(1, 1), bias=False)
      (bn3): BatchNorm2d(256, eps=1e-05, momentum=0.1, affine=True, track_running_stats=True)
      (relu): ReLU(inplace=True)
      (downsample): Sequential(
        (0): Conv2d(64, 256, kernel_size=(1, 1), stride=(1, 1), bias=False)
        (1): BatchNorm2d(256, eps=1e-05, momentum=0.1, affine=True, track_running_stats=True)
      )
    )
    (1): Bottleneck(
      (conv1): Conv2d(256, 64, kernel_size=(1, 1), stride=(1, 1), bias=False)
      (bn1): BatchNorm2d(64, eps=1e-05, momentum=0.1, affine=True, track_running_stats=True)
      (conv2): Conv2d(64, 64, kernel_size=(3, 3), stride=(1, 1), padding=(1, 1), bias=False)
      (bn2): BatchNorm2d(64, eps=1e-05, momentum=0.1, affine=True, track_running_stats=True)
      (conv3): Conv2d(64, 256, kernel_size=(1, 1), stride=(1, 1), bias=False)
      (bn3): BatchNorm2d(256, eps=1e-05, momentum=0.1, affine=True, track_running_stats=True)
      (relu): ReLU(inplace=True)
    )
    (2): Bottleneck(
      (conv1): Conv2d(256, 64, kernel_size=(1, 1), stride=(1, 1), bias=False)
      (bn1): BatchNorm2d(64, eps=1e-05, momentum=0.1, affine=True, track_running_stats=True)
      (conv2): Conv2d(64, 64, kernel_size=(3, 3), stride=(1, 1), padding=(1, 1), bias=False)
      (bn2): BatchNorm2d(64, eps=1e-05, momentum=0.1, affine=True, track_running_stats=True)
      (conv3): Conv2d(64, 256, kernel_size=(1, 1), stride=(1, 1), bias=False)
      (bn3): BatchNorm2d(256, eps=1e-05, momentum=0.1, affine=True, track_running_stats=True)
      (relu): ReLU(inplace=True)
    )
  )
  (layer2): Sequential(
    (0): Bottleneck(
      (conv1): Conv2d(256, 128, kernel_size=(1, 1), stride=(1, 1), bias=False)
      (bn1): BatchNorm2d(128, eps=1e-05, momentum=0.1, affine=True, track_running_stats=True)
      (conv2): Conv2d(128, 128, kernel_size=(3, 3), stride=(2, 2), padding=(1, 1), bias=False)
      (bn2): BatchNorm2d(128, eps=1e-05, momentum=0.1, affine=True, track_running_stats=True)
      (conv3): Conv2d(128, 512, kernel_size=(1, 1), stride=(1, 1), bias=False)
      (bn3): BatchNorm2d(512, eps=1e-05, momentum=0.1, affine=True, track_running_stats=True)
      (relu): ReLU(inplace=True)
      (downsample): Sequential(
        (0): Conv2d(256, 512, kernel_size=(1, 1), stride=(2, 2), bias=False)
        (1): BatchNorm2d(512, eps=1e-05, momentum=0.1, affine=True, track_running_stats=True)
      )
    )
    (1): Bottleneck(
      (conv1): Conv2d(512, 128, kernel_size=(1, 1), stride=(1, 1), bias=False)
      (bn1): BatchNorm2d(128, eps=1e-05, momentum=0.1, affine=True, track_running_stats=True)
      (conv2): Conv2d(128, 128, kernel_size=(3, 3), stride=(1, 1), padding=(1, 1), bias=False)
      (bn2): BatchNorm2d(128, eps=1e-05, momentum=0.1, affine=True, track_running_stats=True)
      (conv3): Conv2d(128, 512, kernel_size=(1, 1), stride=(1, 1), bias=False)
      (bn3): BatchNorm2d(512, eps=1e-05, momentum=0.1, affine=True, track_running_stats=True)
      (relu): ReLU(inplace=True)
    )
    (2): Bottleneck(
      (conv1): Conv2d(512, 128, kernel_size=(1, 1), stride=(1, 1), bias=False)
      (bn1): BatchNorm2d(128, eps=1e-05, momentum=0.1, affine=True, track_running_stats=True)
      (conv2): Conv2d(128, 128, kernel_size=(3, 3), stride=(1, 1), padding=(1, 1), bias=False)
      (bn2): BatchNorm2d(128, eps=1e-05, momentum=0.1, affine=True, track_running_stats=True)
      (conv3): Conv2d(128, 512, kernel_size=(1, 1), stride=(1, 1), bias=False)
      (bn3): BatchNorm2d(512, eps=1e-05, momentum=0.1, affine=True, track_running_stats=True)
      (relu): ReLU(inplace=True)
    )
    (3): Bottleneck(
      (conv1): Conv2d(512, 128, kernel_size=(1, 1), stride=(1, 1), bias=False)
      (bn1): BatchNorm2d(128, eps=1e-05, momentum=0.1, affine=True, track_running_stats=True)
      (conv2): Conv2d(128, 128, kernel_size=(3, 3), stride=(1, 1), padding=(1, 1), bias=False)
      (bn2): BatchNorm2d(128, eps=1e-05, momentum=0.1, affine=True, track_running_stats=True)
      (conv3): Conv2d(128, 512, kernel_size=(1, 1), stride=(1, 1), bias=False)
      (bn3): BatchNorm2d(512, eps=1e-05, momentum=0.1, affine=True, track_running_stats=True)
      (relu): ReLU(inplace=True)
    )
  )
  (layer3): Sequential(
    (0): Bottleneck(
      (conv1): Conv2d(512, 256, kernel_size=(1, 1), stride=(1, 1), bias=False)
      (bn1): BatchNorm2d(256, eps=1e-05, momentum=0.1, affine=True, track_running_stats=True)
```

```
      (conv2): Conv2d(256, 256, kernel_size=(3, 3), stride=(2, 2), padding=(1, 1), bias=False)
      (bn2): BatchNorm2d(256, eps=1e-05, momentum=0.1, affine=True, track_running_stats=True)
      (conv3): Conv2d(256, 1024, kernel_size=(1, 1), stride=(1, 1), bias=False)
      (bn3): BatchNorm2d(1024, eps=1e-05, momentum=0.1, affine=True, track_running_stats=True)
      (relu): ReLU(inplace=True)
      (downsample): Sequential(
        (0): Conv2d(512, 1024, kernel_size=(1, 1), stride=(2, 2), bias=False)
        (1): BatchNorm2d(1024, eps=1e-05, momentum=0.1, affine=True, track_running_stats=True)
      )
    )
    (1): Bottleneck(
      (conv1): Conv2d(1024, 256, kernel_size=(1, 1), stride=(1, 1), bias=False)
      (bn1): BatchNorm2d(256, eps=1e-05, momentum=0.1, affine=True, track_running_stats=True)
      (conv2): Conv2d(256, 256, kernel_size=(3, 3), stride=(1, 1), padding=(1, 1), bias=False)
      (bn2): BatchNorm2d(256, eps=1e-05, momentum=0.1, affine=True, track_running_stats=True)
      (conv3): Conv2d(256, 1024, kernel_size=(1, 1), stride=(1, 1), bias=False)
      (bn3): BatchNorm2d(1024, eps=1e-05, momentum=0.1, affine=True, track_running_stats=True)
      (relu): ReLU(inplace=True)
    )
    (2): Bottleneck(
      (conv1): Conv2d(1024, 256, kernel_size=(1, 1), stride=(1, 1), bias=False)
      (bn1): BatchNorm2d(256, eps=1e-05, momentum=0.1, affine=True, track_running_stats=True)
      (conv2): Conv2d(256, 256, kernel_size=(3, 3), stride=(1, 1), padding=(1, 1), bias=False)
      (bn2): BatchNorm2d(256, eps=1e-05, momentum=0.1, affine=True, track_running_stats=True)
      (conv3): Conv2d(256, 1024, kernel_size=(1, 1), stride=(1, 1), bias=False)
      (bn3): BatchNorm2d(1024, eps=1e-05, momentum=0.1, affine=True, track_running_stats=True)
      (relu): ReLU(inplace=True)
    )
    (3): Bottleneck(
      (conv1): Conv2d(1024, 256, kernel_size=(1, 1), stride=(1, 1), bias=False)
      (bn1): BatchNorm2d(256, eps=1e-05, momentum=0.1, affine=True, track_running_stats=True)
      (conv2): Conv2d(256, 256, kernel_size=(3, 3), stride=(1, 1), padding=(1, 1), bias=False)
      (bn2): BatchNorm2d(256, eps=1e-05, momentum=0.1, affine=True, track_running_stats=True)
      (conv3): Conv2d(256, 1024, kernel_size=(1, 1), stride=(1, 1), bias=False)
      (bn3): BatchNorm2d(1024, eps=1e-05, momentum=0.1, affine=True, track_running_stats=True)
      (relu): ReLU(inplace=True)
    )
    (4): Bottleneck(
      (conv1): Conv2d(1024, 256, kernel_size=(1, 1), stride=(1, 1), bias=False)
      (bn1): BatchNorm2d(256, eps=1e-05, momentum=0.1, affine=True, track_running_stats=True)
      (conv2): Conv2d(256, 256, kernel_size=(3, 3), stride=(1, 1), padding=(1, 1), bias=False)
      (bn2): BatchNorm2d(256, eps=1e-05, momentum=0.1, affine=True, track_running_stats=True)
      (conv3): Conv2d(256, 1024, kernel_size=(1, 1), stride=(1, 1), bias=False)
      (bn3): BatchNorm2d(1024, eps=1e-05, momentum=0.1, affine=True, track_running_stats=True)
      (relu): ReLU(inplace=True)
    )
    (5): Bottleneck(
      (conv1): Conv2d(1024, 256, kernel_size=(1, 1), stride=(1, 1), bias=False)
      (bn1): BatchNorm2d(256, eps=1e-05, momentum=0.1, affine=True, track_running_stats=True)
      (conv2): Conv2d(256, 256, kernel_size=(3, 3), stride=(1, 1), padding=(1, 1), bias=False)
      (bn2): BatchNorm2d(256, eps=1e-05, momentum=0.1, affine=True, track_running_stats=True)
      (conv3): Conv2d(256, 1024, kernel_size=(1, 1), stride=(1, 1), bias=False)
      (bn3): BatchNorm2d(1024, eps=1e-05, momentum=0.1, affine=True, track_running_stats=True)
      (relu): ReLU(inplace=True)
    )
  )
  (layer4): Sequential(
    (0): Bottleneck(
      (conv1): Conv2d(1024, 512, kernel_size=(1, 1), stride=(1, 1), bias=False)
      (bn1): BatchNorm2d(512, eps=1e-05, momentum=0.1, affine=True, track_running_stats=True)
      (conv2): Conv2d(512, 512, kernel_size=(3, 3), stride=(2, 2), padding=(1, 1), bias=False)
      (bn2): BatchNorm2d(512, eps=1e-05, momentum=0.1, affine=True, track_running_stats=True)
      (conv3): Conv2d(512, 2048, kernel_size=(1, 1), stride=(1, 1), bias=False)
      (bn3): BatchNorm2d(2048, eps=1e-05, momentum=0.1, affine=True, track_running_stats=True)
      (relu): ReLU(inplace=True)
      (downsample): Sequential(
        (0): Conv2d(1024, 2048, kernel_size=(1, 1), stride=(2, 2), bias=False)
        (1): BatchNorm2d(2048, eps=1e-05, momentum=0.1, affine=True, track_running_stats=True)
      )
    )
    (1): Bottleneck(
      (conv1): Conv2d(2048, 512, kernel_size=(1, 1), stride=(1, 1), bias=False)
      (bn1): BatchNorm2d(512, eps=1e-05, momentum=0.1, affine=True, track_running_stats=True)
      (conv2): Conv2d(512, 512, kernel_size=(3, 3), stride=(1, 1), padding=(1, 1), bias=False)
      (bn2): BatchNorm2d(512, eps=1e-05, momentum=0.1, affine=True, track_running_stats=True)
      (conv3): Conv2d(512, 2048, kernel_size=(1, 1), stride=(1, 1), bias=False)
      (bn3): BatchNorm2d(2048, eps=1e-05, momentum=0.1, affine=True, track_running_stats=True)
      (relu): ReLU(inplace=True)
    )
    (2): Bottleneck(
      (conv1): Conv2d(2048, 512, kernel_size=(1, 1), stride=(1, 1), bias=False)
      (bn1): BatchNorm2d(512, eps=1e-05, momentum=0.1, affine=True, track_running_stats=True)
      (conv2): Conv2d(512, 512, kernel_size=(3, 3), stride=(1, 1), padding=(1, 1), bias=False)
      (bn2): BatchNorm2d(512, eps=1e-05, momentum=0.1, affine=True, track_running_stats=True)
      (conv3): Conv2d(512, 2048, kernel_size=(1, 1), stride=(1, 1), bias=False)
      (bn3): BatchNorm2d(2048, eps=1e-05, momentum=0.1, affine=True, track_running_stats=True)
      (relu): ReLU(inplace=True)
    )
  )
  (avgpool): AdaptiveAvgPool2d(output_size=(1, 1))
  (fc): Linear(in_features=2048, out_features=1000, bias=True)
)
```

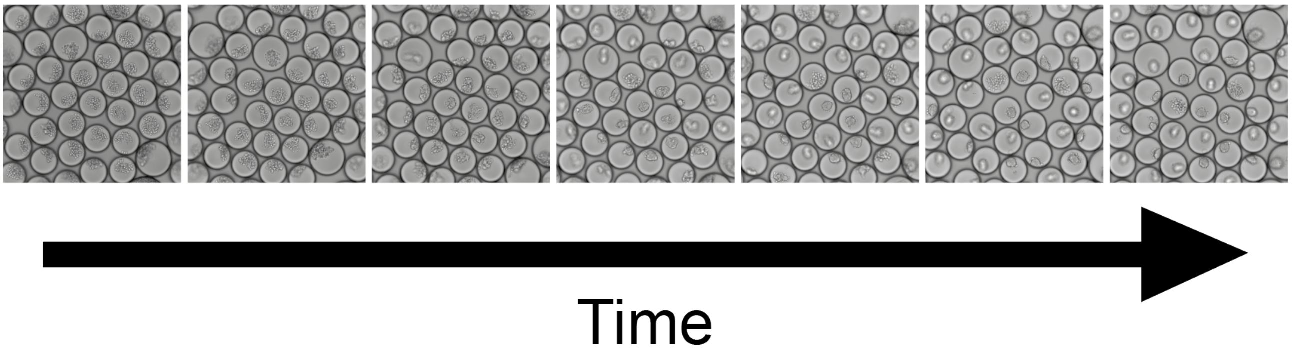


**Supplemental Figure 1.** RT4 cells cultured in Novec HFE-7500 with 0.5 % w/v surfactant, imaged over time. Time points shown are 2 h, 4 h, 6 h, 8 h, 10 h, 12 h and 14 h at the same focal plane. The droplet focal plane remains the same for the duration of the experiment, ruling out evaporation. Some cell ensembles within the droplets move out of focus, though.

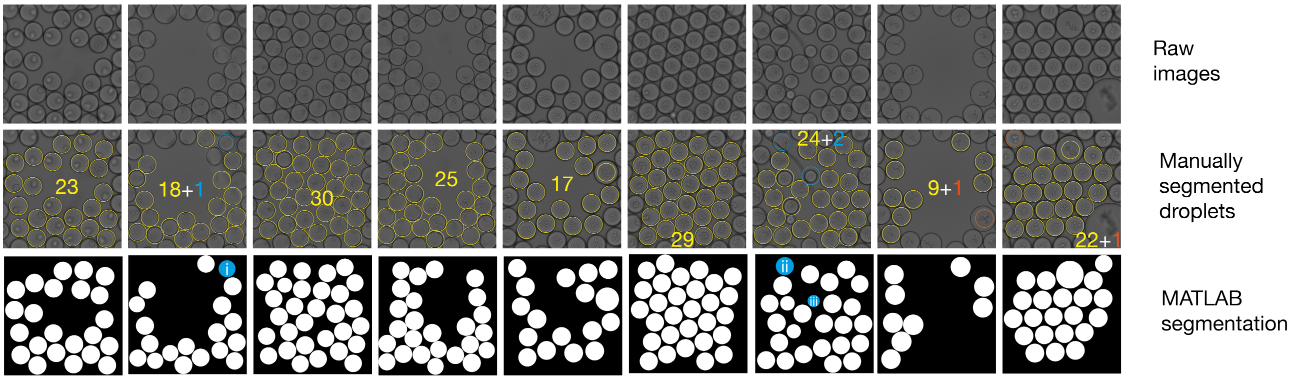


**Supplemental Figure 2.** Segmentation validation. Randomly selected raw images were manually assessed for droplet number and compared to the MATLAB segmentation. Manual segmentation would have resulted in 199 droplets, 197 of which (yellow) were also segmented using the MATLAB script (99 %). The missed droplets are highlighted in orange. The script segmented an additional 3 droplets (blue), which we would not have picked because of an imaging artifact (i), an object obstructing the view (ii) or because it was empty (iii) (1.5 % error rate).

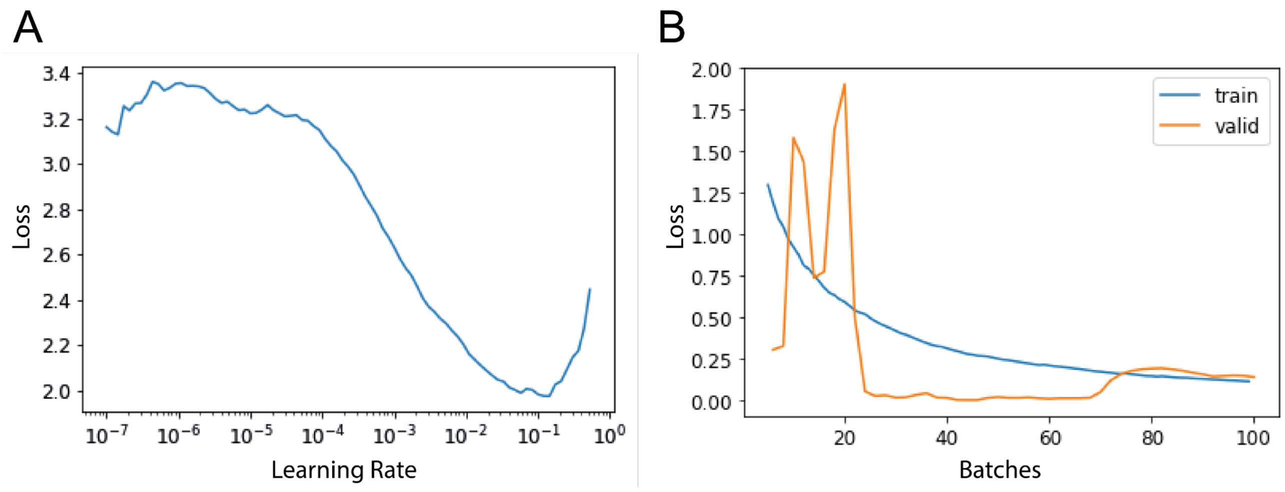


**Supplemental Figure 3.** Learning rate and training plots. A) Learning rate plot for SpheroClassCNN. A safe learning rate is around an order of magnitude lower than the learning rate with the lowest loss before the spike, in our case at 9.12 x $10^{-3}$. B) Training plot for BlurredObjectsCNN. After 20 batches the validation loss sharply decreases (orange line). After roughly 70 batches the validation loss starts to rise again while the training loss (blue line) continues to fall, signifying overfitting.

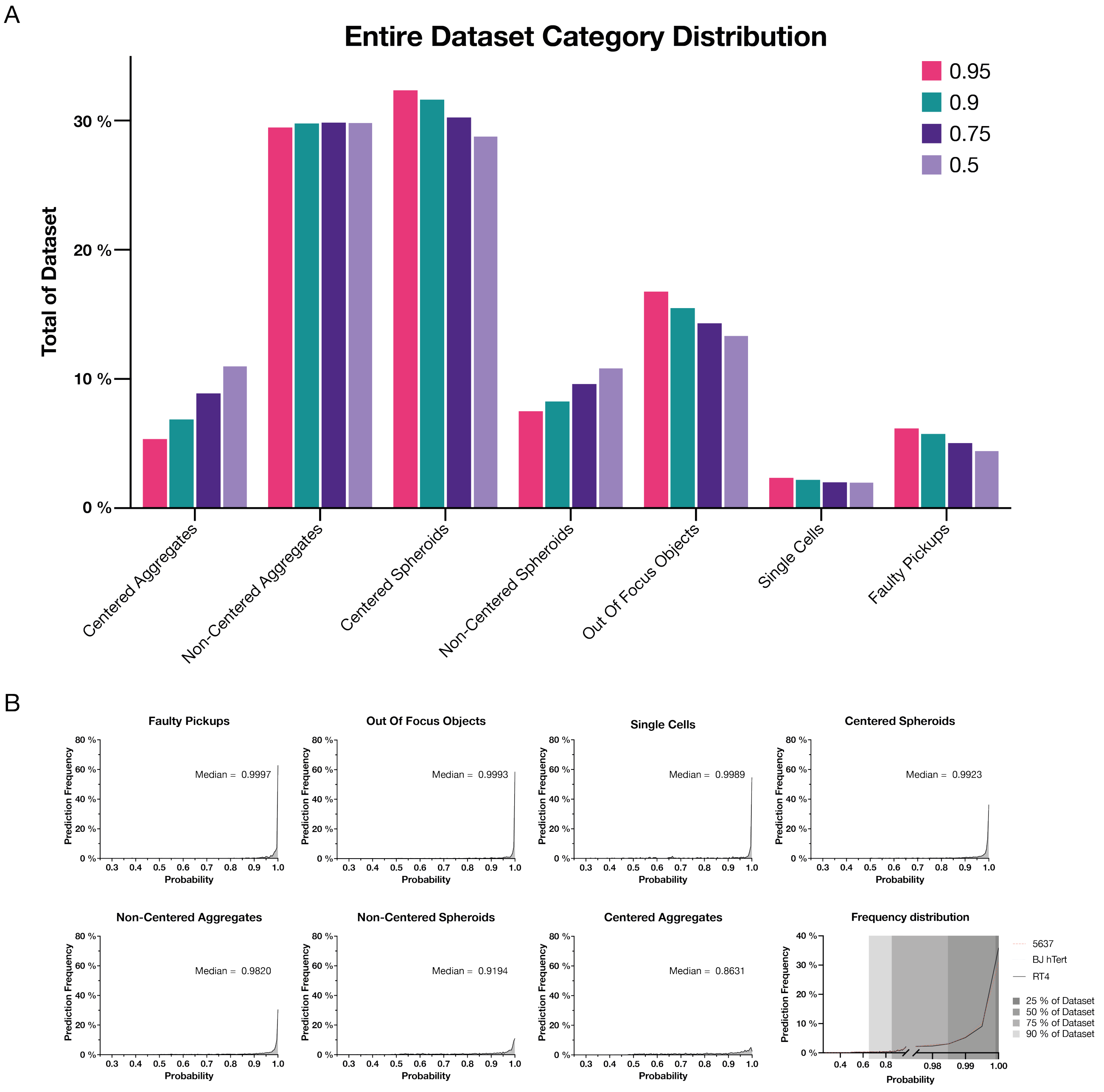


**Supplemental Figure 4.** Category distribution and assigned probability plots. A) shows the distribution of categories of the entire SpheroClassCNN-analyzed dataset with different probability thresholds. B) shows the frequency of assigned probabilities for the different categories and for the different cell lines. Grey rectangles represent percentages of the dataset.

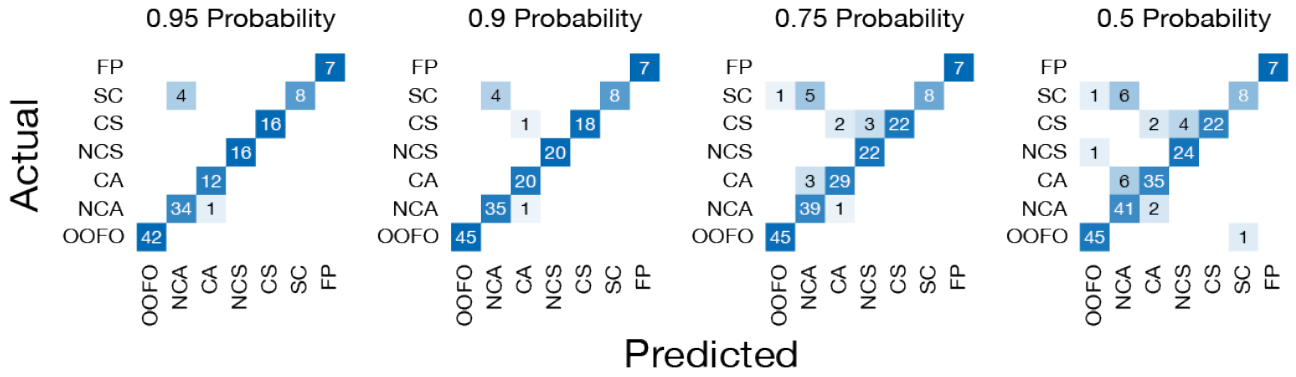


**Supplemental Figure 5.** Confusion matrices showing the CNN (predicted) versus human (actual) categorizations over the different probability thresholds. CA = Centered Aggregate, NCA = Non-Centered Aggregate, CS = Centered Spheroid, NCS = Non-Centered Spheroid, OOFO = Out Of Focus Object, SC = Single Cells, FP = Faulty Pickup.

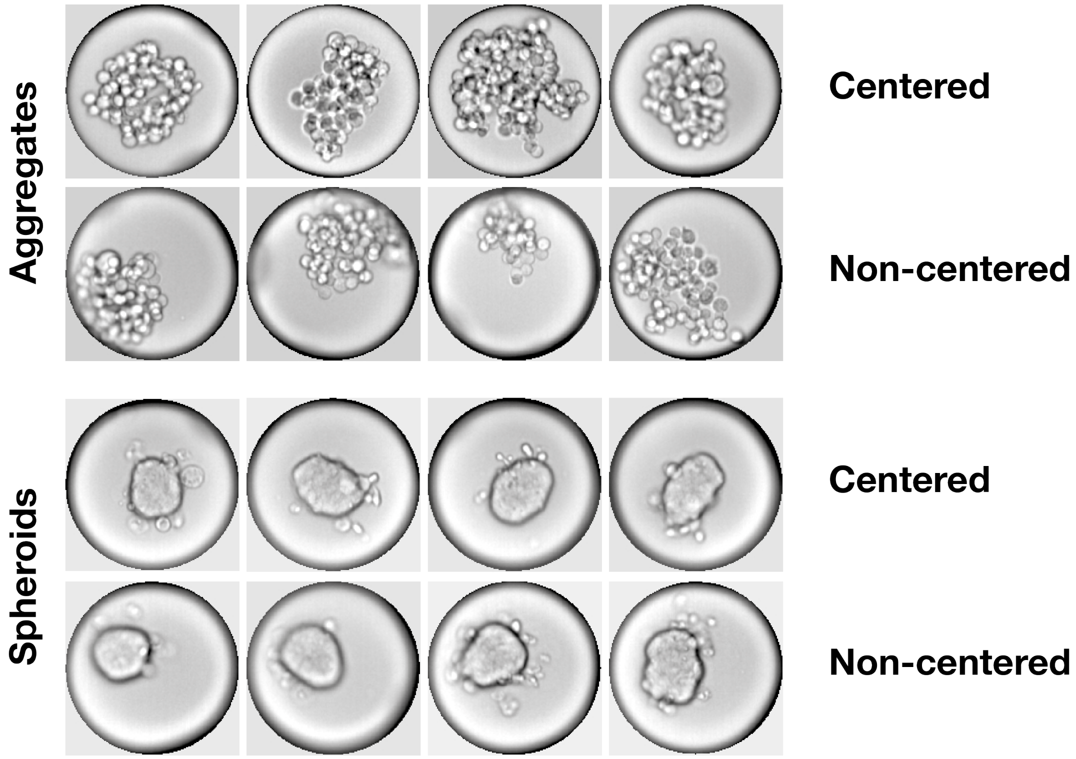


**Supplemental Figure 6.** Exemplary single-droplet images comparing intra-categorical morphological diversity. Spheroids show a characteristically strong contrast at the border of the object, whereas aggregates are inherently more diverse.

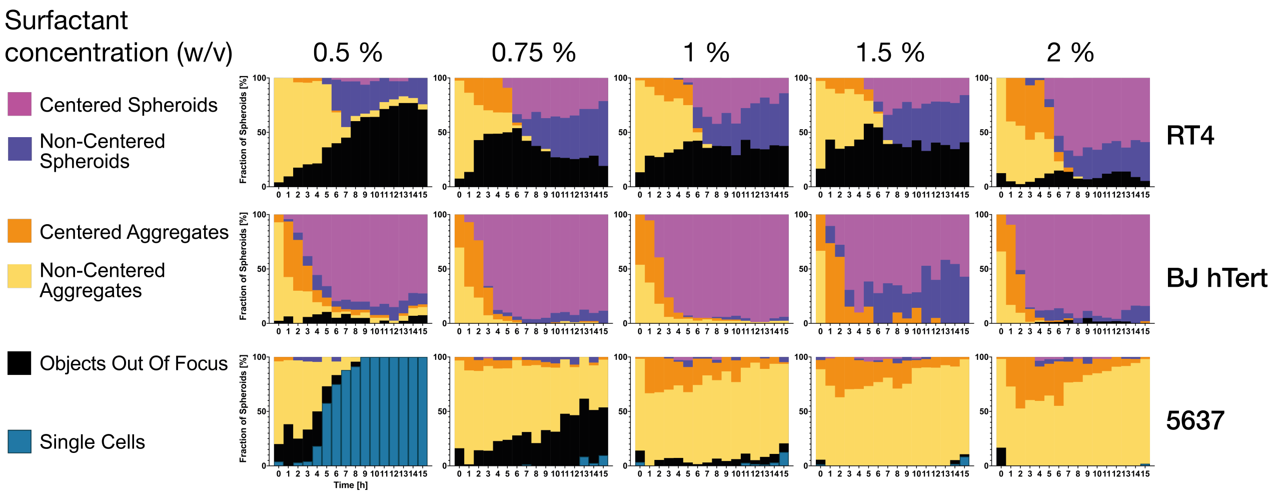


**Supplemental Figure 7.** SpheroClassCNN classification of droplets. The Objects Out Of Focus class shows up in all three cell lines and makes up a considerable fraction in the RT4 sample. We subsequently trained the BlurredObjectsCNN to further resolve morphologies.

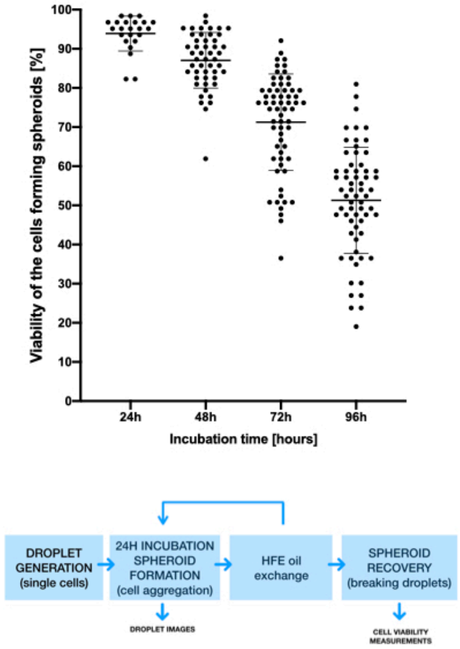


**Supplemental Figure 8.** Viability of HepG2 spheroids over time. HepG2 cells were encapsulated and incubated for up to 96 h. For each time point emulsion was broken and spheroids were stained using Hoechst 33342, Propidium Iodide and Calcein AM.

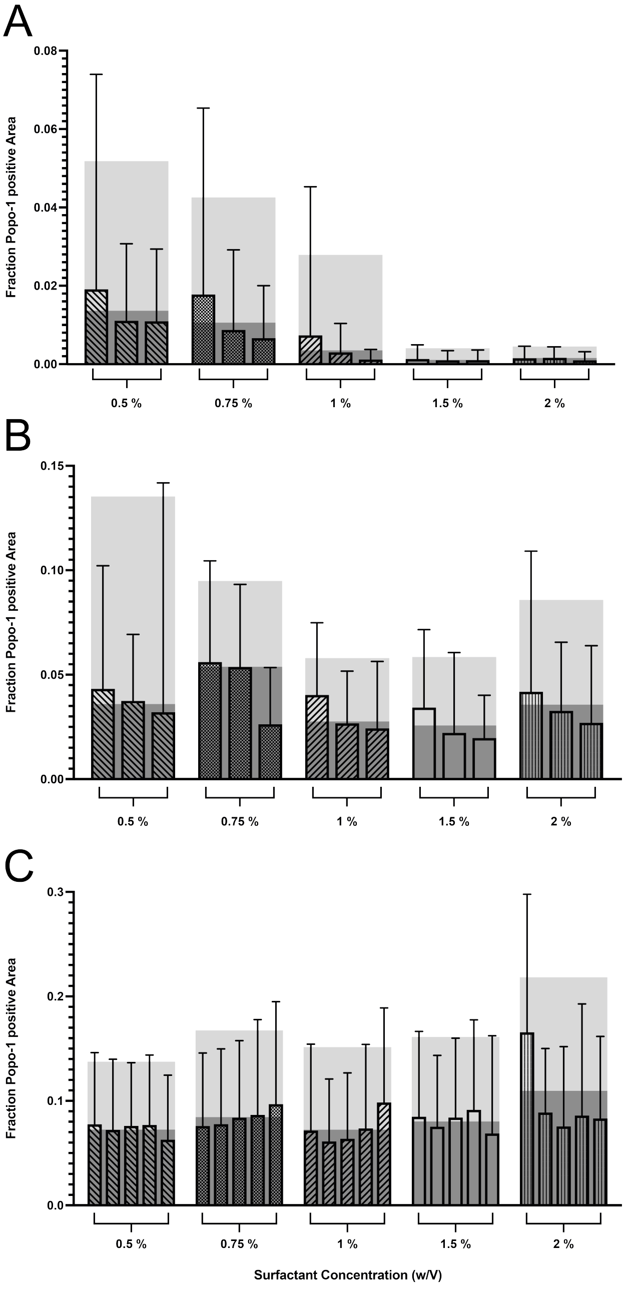


**Supplemental Figure 9.** Popo-1 iodide fluorescence analysis after 15 hours. Patterned bars show the results of individual wells, dark grey boxes illustrate the mean for the respective surfactant concentration, light grey boxes the error. 5637 cells show a tenfold decrease of Popo-1 iodide-positive area from 0.5 % to 2 % surfactant concentration (n=1152). RT4 (n=955) and BJ hTert cells (n=1847) seem not to be affected in a similar way.

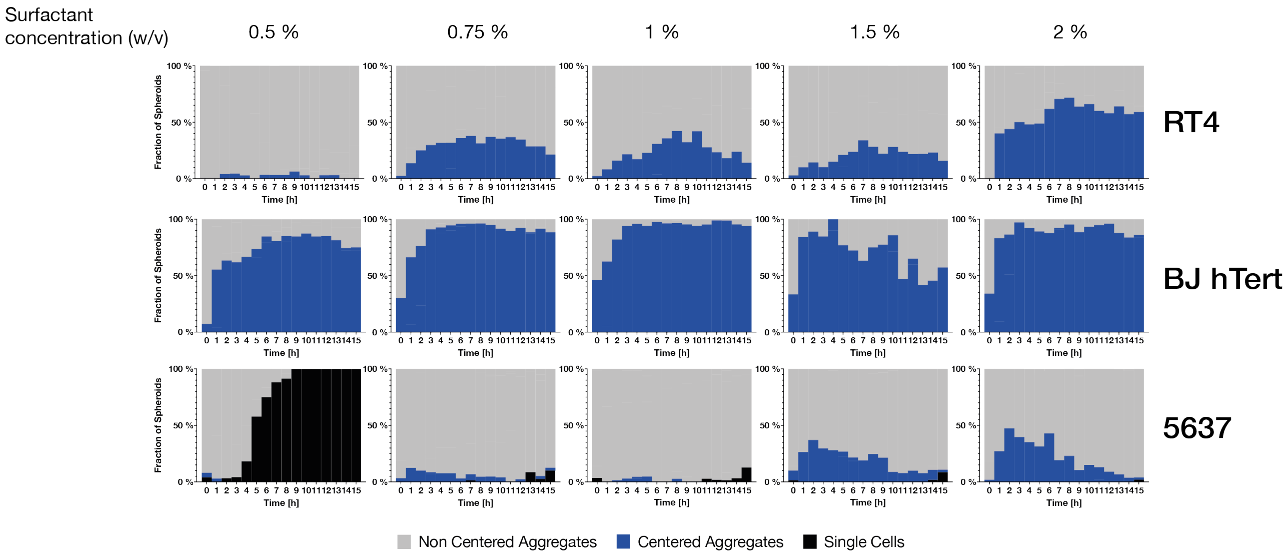


**Supplemental Figure 10.** Droplet classification results broken down according to positioning. We posit that any diversion from the x, y, z center is a consequence of interface interaction. Notably, this also seems to correlate with the individual cell line's spheroid formation capabilities.

### Scripts

The MATLAB and Python scripts developed for this study are available at https://github.com/Schappison/Spheroid-Classifier.
